# Anisotropic mineralized collagen scaffolds accelerate osteogenic response in a glycosaminoglycan-dependent fashion

**DOI:** 10.1101/2020.01.19.911305

**Authors:** Marley J. Dewey, Andrey V. Nosatov, Kiran Subedi, Brendan Harley

## Abstract

Regeneration of critically-sized craniofacial bone defects requires a template to promote cell activity and bone remodeling. However, induced regeneration becomes more challenging with increasing defect size. Methods of repair using allografts and autografts have inconsistent results, attributed to age-related regenerative capabilities of bone. We are developing a mineralized collagen scaffold to promote craniomaxillofacial bone regeneration as an alternative to repair. Here, we hypothesize modifying the pore anisotropy and glycosaminoglycan content of the scaffold will improve cell migration, viability, and subsequent bone formation. Using anisotropic and isotropic scaffold variants, we test the role of pore orientation on human mesenchymal stem cell (MSC) activity. We subsequently explore the role of glycosaminoglycan content, notably chondroitin-6-sulfate, chondroitin-4-sulfate, and heparin sulfate on mineralization. We find that while short term MSC migration and activity was not affected by pore orientation, increased bone mineral synthesis was observed in anisotropic scaffolds. Further, while scaffold glycosaminoglycan content did not impact cell viability, heparin sulfate and chondroitin-6-sulfate containing variants increased mineral formation at the late stage of *in vitro* culture, respectively. Overall, these findings show scaffold microstructural and proteoglycan modifications represent a powerful tool to improve MSC osteogenic activity.

## 1. Introduction

Osseous defects of the skull occur secondary to trauma, congenital abnormalities, or after resection to treat stroke, cerebral aneurysms, or cancer ^1–6^. While bone has the capacity to regenerate over time, the large scale of craniomaxillofacial (CMF) defects typically preclude effective regeneration. Thus, solutions to regenerate bone in these defects involve the use of autografts, allografts, and other biomaterial implants. Autografts and allografts are the typical standard for repair, using bone from a patient’s own body or from a donor. These methods of repair have variable success rates, due to age-related regenerative property changes in bone grafts ^7–9^. Other biomaterial strategies seek to overcome these limitations, and one such material of interest is mineralized collagen. Mineralized collagen scaffolds are composed of the natural components of bone (type I collagen, calcium and phosphate ions, glycosaminoglycans), are highly porous, and have been shown to promote mineral formation *in vitro* without osteogenic supplements and produce bone *in vivo* ^10–19^. The scale of critical sized defects is a challenge to these and other biomaterial approaches, which demand designs to improve cell invasion, proliferation, and mineral biosynthesis activity. Current clinical strategies often use large exogenous doses of growth factors (*e*.*g*., bone morphogenic protein 2, BMP-2) ^18^, that have significant complications such as ectopic bone formation, resorption of adjacent bone, and maxillary growth disorders ^20–24^. These drawbacks highlight the need for a new paradigm.

We are developing a material-only biomaterial approach to actively promote, rather than passively permit, MSC recruitment, osteogenic differentiation, and eventually bone regeneration. Pore size and architecture remain an important factor in guiding tissue regeneration. Pore sizes in the range of 100-350 μm have been shown to promote the formation of bone ^25^. Further, the native anisotropy of trabecular bone is increasingly being explored as a design paradigm ^26^. Recently, Klaumünzer *et al*., suggested anisotropic pores in a collagen biomaterial can better promote intramembranous ossification required in CMF bone repair ^26^. We have previously developed an approach to fabricate non-mineralized collagen scaffolds with anisotropic pores to increase the metabolic activity of tenocytes and provide directional growth of these cells for tendon repair ^27–31^. Further, glycosaminoglycans (GAGs) in the extracellular matrix play an important role in bone homeostasis and potentially regeneration ^32^. GAGs linked to bone repair include chondroitin-6-sulfate (CS6), chondroitin-4-sulfate (CS4), and heparin sulfate (Heparin). CS6 has been shown to be the main GAG present in articular cartilage, while CS4 is hypothesized to be more important to calcification ^33^ and collagen-CS4 biomaterials can promote *in vitro* bone mineralization ^34^. Alternatively, non-covalent interactions with sulfated GAGs such as heparin sulfate can enhance retention and activity of osteogenic growth factors such as BMP-2 to promote osteogenic activity ^35–38^. We know the GAG chemistry of non-mineralized collagen scaffolds can be exploited to enhance growth factor sequestration and resultant cell activity ^39^.

The mineralized collagen scaffolds under development in our lab possess isotropic pores and contain CS6 glycosaminoglycans. In this work we seek to modify the pore anisotropy and GAG content of these scaffolds to increase MSC invasion and osteogenic activity. We hypothesize: (1) anisotropic pores within the mineralized scaffolds will increase cell invasion and osteogenic activity; and, (2) substituting heparin sulfate or chondroitin-4-sulfate for chondroitin-6-sulfate in these scaffolds will increase bone formation. We report the porosity and pore orientation of a family of anisotropic and isotropic scaffolds, examine MSC invasion into the scaffold using a modified transwell membrane assay, and evaluate human MSC osteogenic activity and mineral biosynthesis over 28 days of *in vitro* culture.

## 2. Materials and Methods

### 2.1. Experimental Design

The aim of this study was to determine mineral formation and cellular activity changes of human mesenchymal stem cells (hMSCs) seeded on mineralized collagen scaffolds as a function of pore orientation and glycosaminoglycan content. Firstly, we compared cell invasion into, subsequent bioactivity, and mineral formation as a function of pore anisotropy for a family of mineralized collagen scaffolds with a conventional glycosaminoglycan content (chondroitin-6-sulfate). Secondly, we compared cell activity and mineral formation of mineralized collagen scaffolds with anisotropic pores as a function of glycosaminoglycan content (chondroitin-6-sulfate, chondroitin-4-sulfate, heparin sulfate). All comparisons involved testing cell viability, gene expression, cytokine release, and mineral formation via alkaline phosphatase, microcomputed tomography (Micro-CT), and Inductively Coupled Plasma (ICP) assays (**Fig. 1**).

**Figure 1:**
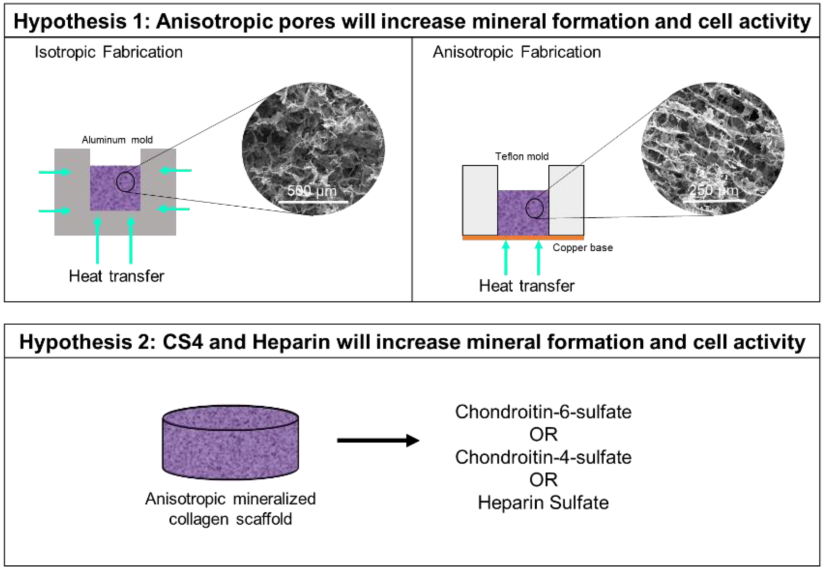
Experimental Outline. Hypothesis 1: Anisotropic pores will have greater mineral formation and cell activity than isotropic pores. The different molds used during freeze drying to fabricate isotropic and anisotropic mineralized collagen scaffolds is demonstrated. Hypothesis 2: Chondroitin-4-sulfate and heparin sulfate will have greater mineral formation and cell activity than scaffolds containing chondroitin-6-sulfate. Anisotropic scaffolds containing the same amount of chondroitin-6-sulfate, chondroitin-4-sulfate, or heparin sulfate are compared.

### 2.2. Fabrication of isotropic and anisotropic mineralized collagen scaffolds

Mineralized collagen scaffolds were fabricated using previously described procedures^10,11,21,27,40–45^. In a cooled, jacketed vessel, 1.9 w/v% type I bovine collagen (Sigma-Aldrich, Missouri, USA), 0.84 w/v% chondroitin-6-sulfate (Sigma-Aldrich), and Ca(OH)2, H3PO4, and Ca(NO3)2·4H2O were thoroughly homogenized, making sure to prevent collagen clumping. The mineralized collagen suspension was transferred to 7.62 × 7.62 cm aluminum molds to create isotropic scaffolds. Alternatively, the suspension was pipetted into Teflon molds with a copper base to create anisotropic scaffolds ^46^. The suspensions were cooled at a constant rate of 1°C/min from 20°C to -10°C. Once frozen, the suspension was held at this temperature for 2 hours then lyophilized (0.2 Torr pressure, 0°C) in a VirTis Genesis 25XL Lyophilizer (SP Industries, Inc., Pennsylvania, USA).

Anisotropic mineralized collagen scaffolds with disparate GAG content were fabricated following the same method. However initial mineralized collagen suspensions were created using one of three potential glycosaminoglycans: chondroitin-6-sulfate (Chondroitin sulfate sodium salt from shark cartilage, Sigma-Aldrich), chondroitin-4-sulfate (Sodium Chondroitin Sulfate A, Toronto Research Chemicals Inc., Ontario, Canada), or heparin sulfate (Heparin sodium salt from porcine intestinal mucosa, Sigma-Aldrich).

### 2.3. SEM imaging and pore size analysis of mineralized collagen scaffolds

A scanning electron microscope (SEM) was used to visualize pore orientation and mineral crystals in isotropic, anisotropic, and various glycosaminoglycan-containing mineralized collagen scaffolds. Scaffolds were cut in half to expose the interior, and then sputter coated with Au/Pd prior to visualizing with a FEI Quanta FEG 450 ESEM (FEI, Oregon, USA). The pore size of isotropic and anisotropic mineralized collagen scaffolds was analyzed following a JB-4 (Polysciences, Inc., Pennsylvania, USA) embedding procedure ^30,47^. After hydrating scaffolds in ethanol and subsequently JB-4, the scaffolds were placed into wells with three samples of each type placed flat into the mold and three placed on their side to create transverse and longitudinal sections. Polymerization was completed overnight at 4°C. Embedded scaffolds were further embedded in paraffin to fit molds for microtome sectioning, and 5 µm sections were cut using a RM2255 microtome (Leica, Wetzlar, Germany) with a tungsten carbide blade. Sectioned scaffolds were placed onto glass slides and stained with an aniline blue solution (Thermo Fisher Scientific, Massachusetts, USA) to stain collagen fibers. Slides were then imaged with a NanoZoomer Digital Pathology System (Hamamatsu, Japan) and pore structure was analyzed with a custom Matlab pore size code to acquire an average pore size and pore aspect ratio for each scaffold ^30,41^.

### 2.4. Sterilization, hydration, and crosslinking of mineralized collagen scaffolds

All scaffolds were sterilized using a dehydrothermal crosslinking and sterilization procedure ^48^. Scaffolds were placed under vacuum in a vacuum oven set at 105°C for 24hrs to complete crosslinking and sterilization ^42^. All subsequent methods followed sterile techniques. Scaffolds were hydrated and further crosslinked before cell seeding. Briefly, scaffolds were soaked in 100% ethanol, followed by multiple washes in PBS, and then crosslinked with EDC-NHS carbodiimide chemistry. Scaffolds were then washed in Phosphate Buffered Saline (PBS) and soaked in normal growth media for 48 hours before cell seeding.

### 2.5. hMSC cell culture and scaffold seeding

Passage 4 human mesenchymal stem cells (Lonza, Switzerland) were expanded to passage 5 in an incubator set at 37°C and 5% CO2 and fed with low glucose DMEM and glutamine, 10% mesenchymal stem cell fetal bovine serum (Gemini, California, USA), and 1% antibiotic-antimycotic (Gibco, Massachusetts, USA). Before seeding cells onto scaffolds, mycoplasma contamination was tested with a MycoAlert™ Mycoplasma Detection Kit (Lonza); all experiments used cells that tested negative for mycoplasma. hMSCS were seeded on hydrated scaffolds at a density of 150,000 cells per scaffold, with 7,500 cells/µL on the bottom and top of each scaffold using a previously defined static seeding method ^47,49^. Seeded scaffolds were placed in an incubator for an hour and a half to allow cell attachment before basal media was added without osteogenic supplements. Media was replaced every third day and the study proceeded for 28 days.

### 2.6. Metabolic activity quantification of cell-seeded scaffolds

An alamarBlue^®^ assay was used to quantify the metabolic activity of all scaffold variants ^27^. A standard curve was generated with known cell numbers to correlate fluorescent readings with fold change of metabolic activity. In reported data, a metabolic activity of 1 represents the metabolic activity of the initial scaffold cell seeding density (150,000 cells). Scaffolds were washed in PBS at every timepoint (day 1, 4, 7, 14, 28) and placed in a solution of alamarBlue^®^ (Invitrogen, California, USA) in an incubator at 37°C on a shaker. After incubation for 1.5 hours, the alamarBlue^®^ solution was collected and fluorescence of resorufin (540(52) nm excitation, 580(20) nm emission) was measured with a F200 spectrophotometer (Tecan, Switzerland) (n=6).

### 2.7. Gene expression quantification of cell-seeded scaffolds

Osteogenic (*RUNX2, COL1A2, Osterix, BMP2)*, chondrogenic (*SOX9*), and adipogenic (*PPARγ*) gene expression was monitored over a 28 day in vitro experiment ^30^. RNA was isolated from cell-seeded scaffolds using a RNeasy Plant Mini Kit (Qiagen, California, USA). RNA was reverse transcribed to cDNA using a QuantiTect Reverse Transcription kit (Qiagen) and a S100 thermal cycler (Bio-Rad, California, USA). 10 ng of cDNA was used in each well of a real-time PCR reaction, using Taqman primers and a Taqman fast advanced master mix (Applied Biosystems, California, USA). Primers can be found listed in **Table 1**. PCR plates were read using a QuantstudioTM 7 Flex Real-Tim PCR System (Thermo Fisher Scientific). Results (n=5) were analyzed using a delta-delta CT method and expressed as fold change normalized to expression levels prior to seeding on scaffolds. *GAPDH* was used as a housekeeping gene.

**Table 1.**
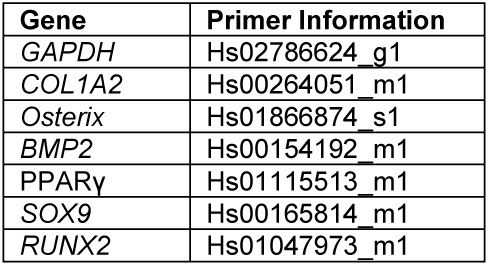
Primers used for RT-PCR.

| Gene | Primer Information |
| --- | --- |
| *GAPDH* | Hs02786624_g1 |
| *COL1A2* | Hs00264051_m1 |
| *Osterix* | Hs01866874_s1 |
| *BMP2* | Hs00154192_m1 |
| *PPARγ* | Hs01115513_m1 |
| *SOX9* | Hs00165814_m1 |
| *RUNX2* | Hs01047973_m1 |

### 2.8. Mineral formation analysis via Micro-CT and ICP Optical Emission Spectrometry

Final mineral formation in cell-seeded scaffolds was measured by Micro-CT and ICP. For Micro-CT analysis, scaffolds were fixed in formalin and then stored in PBS at 4°C prior to imaging. A MicroXCT-400 (Zeiss, Germany) was used to scan samples (n=3) using a 1x magnitude camera and a power set to 8 W and 40kV, with the same X-ray source and detector distance, exposure time, and binning for each scaffold. ImageJ was used to evaluate the mineral content from z-stacks of 2D images, and a threshold intensity of 214/255 was set for every scaffold. The particle average intensity was then calculated in ImageJ, following a previously described procedure ^28^. To calculate average fill of the scaffolds, the average total area of the particles was divided by the area of the scaffold. Scaffold average fill at day 28 was compared to unseeded scaffolds at day 0.

After Micro-CT analysis, scaffolds were analyzed by ICP Optical Emission Spectrometry to define the calcium and phosphorous content ^43^. Samples (n=3) were soaked in 100% ethanol dried with an Autosamdri 931 critical point dryer (Tousimis, Maryland, USA). 0.5 mL to 1 mL of liquid samples and ∼5 mg of solid samples were used for elemental analysis. After carefully weighing/pipetting the samples, they were transferred to digestion tubes, and were further treated with concentrated trace metal grade nitric acid (67-70%) purchased from Fisher Scientific. This was followed by automated sequential microwave digestion in a CEM Mars 6 microwave digester (CEM Microwave Technology Ltd., North Carolina, USA). The final digested product was a transparent, aqueous solution which was further diluted to a volume of 50 mL using DI water, so as to make the final concentration of the acid < 5%. Matrix matched standards were prepared for the calibration curves. The solution was then introduced to inductively coupled plasma-mass spectrometer (ICP-OES, Optima 8300, PerkinElmer, USA) for the elemental analysis in axial mode. Digestion and ICP-MS analysis parameters are listed in **Supplementary Tables 1 and 2**.

### 2.9. Alkaline Phosphatase Activity

A colorimetric Alkaline Phosphatase Assay Kit (Abcam, United Kingdom) was used to measure changes in cell-dependent active bone formation in hMSC-seeded scaffolds as a function of pore orientation and glycosaminoglycan content over 28 days *in vitro* ^50^. Day 0 analysis was performed on 2 unseeded scaffolds of each type; subsequent (day 7, 28) analyses were performed on 2 unseeded and 6 MSC-seeded scaffolds of each type. Scaffolds were washed with PBS, collected in tubes, frozen in liquid nitrogen and stored in a -80° freezer. Scaffolds were homogenized and centrifuged and 70 µL of sample per scaffold was used in conjunction with assay buffer and p-nitrophenylphosphate (pNPP) solution for the incubation; after the incubation period, absorbance was measured with a Tecan Infinite M200 Pro (Tecan) at 405nm. Absorbance values for unseeded day 7 and day 28 scaffolds were subtracted from values for seeded scaffolds of the respective groups, and the normalized values were divided by group-averaged values from day 0 to give a fold change for each scaffold type (n=6).

### 2.10. Cell migration assay using transwell membranes

Cell migration in anisotropic and isotropic scaffolds containing chondroitin-6-sulfate was evaluated via a modified transwell membrane invasion assay (**Supp. Fig. 1**) ^28^. Briefly, hMSCs were seeded on the top surface of 8 µm pore size adjustable transwell membranes (ThermoFisher Scientific). Scaffolds were added to the bottom of the plate with 600 μL of normal growth media containing 50 ng/mL of PDGF-BB (ThermoFisher) to induce chemotactic migration ^30^. Transwells were adjusted to rest on the top of the scaffolds and 50,000 hMSCs were seeded onto the transwell membrane in 100 μL of normal growth media. Scaffolds were allowed to sit for 24hrs, and after 24hrs scaffolds were removed and an alamarBlue^®^ assay was used to quantify the number of cells that had migrated into each scaffold type following the procedure outlined in 2.6 and a standard curve using 50,000 cells.

### 2.11. Cytokine release

A RayBio^®^ C-Series Human Cytokine Antibody Array C5 (Ray Biotech, Georgia, USA) was used to measure the relative levels of 80 cytokines eluded from hMSC-seeded scaffolds as a function of pore orientation and glycosaminoglycan content. MSC-seeded scaffolds were incubated for 7 days and media was collected and replaced on the 3^rd^, 6^th^, and 7^th^ days. Following collection, media was pooled for analysis. Cell media not exposed to scaffolds was used as a control. The procedure for the assay was performed in triplicate with 1 mL of undiluted sample for each group. Membranes were imaged with an ImageQuant LAS 4010 (GE Healthcare Life Sciences, Massachusetts, USA) for 1 second. Resulting images were analyzed with ImageJ to measure intensity levels of each cytokine; once measurements were obtained one sample was arbitrarily chosen for positive control normalization. Results were then normalized to the media control by dividing to give a fold increase over the control (n=3).

### 2.12. Statistical Analysis

Statistics were performed with an OriginPro software (OriginPro, Massachusetts, USA). Significance set to p < 0.05 and tested in accordance with literature ^51^. Firstly, a Shapiro-Wilk test was used to test for normality and then a Grubbs outlier test was used to remove any outliers in non-normal data sets. If data still did not meet the normality assumption, a Kruskal-Wallis test was used to evaluate significance in data. For normal data, a Browne-Forsythe test was used to test the equal variance assumption. A t-test with a Welch correction was used between two samples if the equal-variance assumption was not met. A t-test was used for all analyses between only two samples. If the assumption of normality and equal variance was met, then an ANOVA with a Tukey post-hoc test was used to determine significance. For powers below 0.8, the data was deemed inconclusive. Sample numbers used for each group was based on previous studies utilizing similar collagen scaffolds ^27,41–43,52^: metabolic activity (n=6), gene expression (n=5), micro-CT (n=3), ICP (n=3), ALP (n=6), cytokine array (n=3). Error bars for all data are represented as mean ± standard deviation.

## 3. Results

### 3.1. SEM images and pore size analysis demonstrate anisotropic structure of scaffolds

SEM imaging revealed anisotropic scaffolds fabricated via a directional solidification approach demonstrated elongated pores with alignment in one direction (**Supp. Fig. 2**). Stereology was subsequently used to quantify pore size and aspect ratio, revealing a significantly smaller aspect ratio in the transverse (vs. longitudinal) plane of anisotropic scaffolds to confirm structural anisotropy (**Supp. Fig. 3, Table 2**). Averaged across all planes of analysis, there was no significant difference in the mean pore size of anisotropic and isotropic scaffolds.

**Table 2.** Pore size and aspect ratio analysis of chondroitin-6-sulfate isotropic and anisotropic scaffolds cut in the longitudinal and transverse direction. There was no significant difference (p < 0.05) in pore size, however, the pore aspect ratio of the anisotropic scaffolds cut transversely was significantly (p < 0.05) different than the isotropic scaffolds cut longitudinally.

| Pore alignment | Orientation | Average pore aspect ratio | Average pore diameter ($\mu\text{m}$) |
| --- | --- | --- | --- |
| Isotropic | Transverse | $0.948 \pm 0.004$ | $177 \pm 19.3$ |
| | Longitudinal | $0.955 \pm 0.003$ | $169 \pm 5.2$ |
| Anisotropic | Transverse | $0.887 \pm 0.034$ | $170 \pm 25.7$ |
| | Longitudinal | $0.938 \pm 0.007$ | $140 \pm 9.5$ |

### 3.2. All scaffold variants supported increasing MSC metabolic activity over 28 days

Pore orientation did not affect the metabolic activity of hMSCs seeded on scaffolds for 28 days. Isotropic and anisotropic variants both displayed an overall increase in MSC metabolic activity over 28 days in culture (**Fig. 2A**). Changing the glycosaminoglycan type within scaffolds also had no effect on metabolic activity, and at day 28 the metabolic activity of all scaffolds was greater than day 1 of the study (**Fig. 2B**).

**Figure 2:**
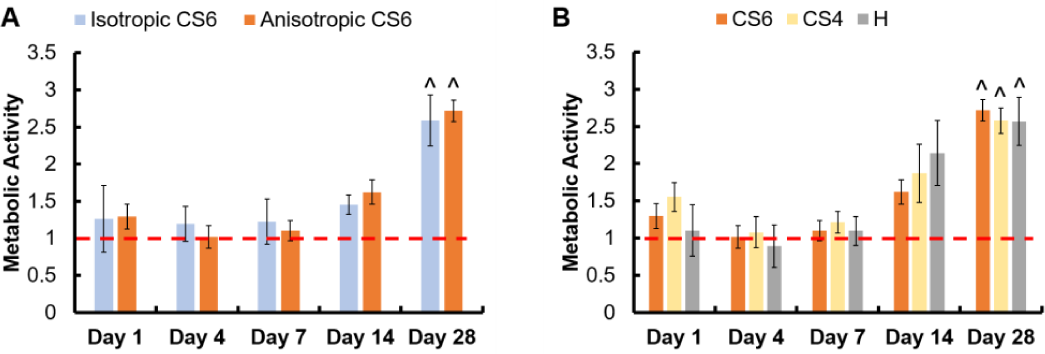
Metabolic activity of hMSC seeded on mineralized collagen scaffolds with isotropic and anisotropic pores and GAGs chondroitin-6-sulfate (CS6), chondroitin-4-sulfate (CS4), and heparin sulfate (H). Metabolic activity was normalized to the metabolic activity of 150,000 hMSCs (cell seeding activity), indicated by a red dashed line. ^ indicates one group was significantly (p < 0.05) greater than the same group on day 1. Data expressed as average ± standard deviation (n=6).

### 3.3. Anisotropic scaffolds had an increase in chondrogenic gene, SOX9, at day 7, but no difference in osteogenic gene expression compared to isotropic variants

Osteogenic genes (*RUNX2, Osterix, BMP2*, and *COL1A2*), chondrogenic gene (*SOX9*), and adipogenic gene (*PPARγ*) were examined in isotropic and anisotropic scaffolds over the course of 28 days. There was an increase in *PPARγ* over 28 days in both scaffold groups, and a spike in *SOX9* expression at day 7 in anisotropic scaffolds (**Fig. 3**). There were no significant differences in all osteogenic gene fold changes between isotropic and anisotropic variants. However, anisotropic scaffolds had an overall increase in gene expression of *COL1A2* by day 7 not seen in isotropic scaffolds.

**Figure 3:**
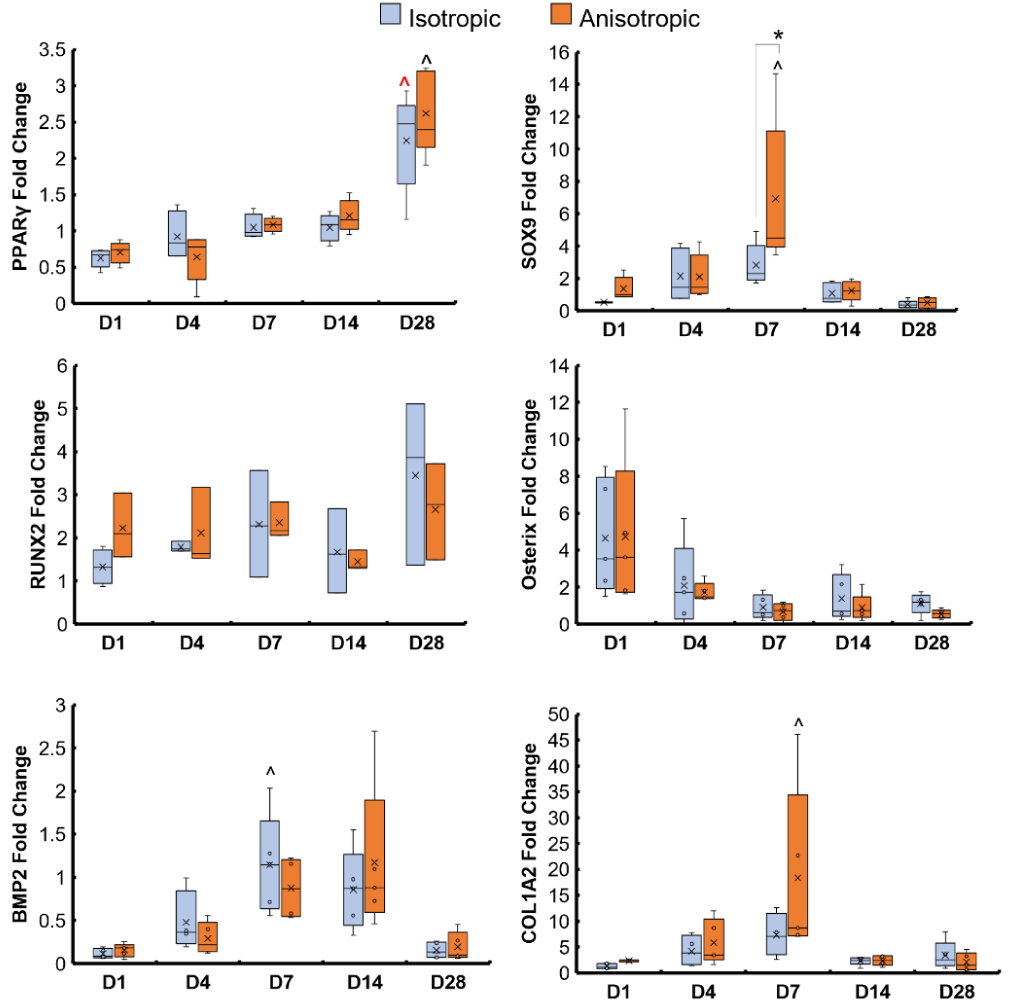
Osteogenic, chondrogenic, and adipogenic gene expression of mineralized collagen scaffolds with anisotropic and isotropic pores. * indicates one group is significantly (p < 0.05) greater than another group on the same day. ^ indicates one group was significantly (p < 0.05) greater than the same group on day 1. Anisotropic scaffolds had significantly (p < 0.05) greater SOX9, chondrogenic, gene expression on day 7 than isotropic scaffolds. Data expressed as average ± standard deviation (n=5).

### 3.4. CS6-containing scaffolds had an increase in chondrogenic gene, SOX9, at day 7, and heparin-containing scaffolds had an increase in Osterix expression at day 4

Osteogenic genes (*RUNX2, Osterix, BMP2*, and *COL1A2*), chondrogenic gene (*SOX9*), and adipogenic gene (*PPARγ*) were examined in scaffolds containing CS6, CS4, and heparin over the course of 28 days (**Fig. 4**). There was an increase in *PPARγ* over 28 days in all scaffold groups, and CS6-containing scaffolds had a greater *SOX9* expression than heparin-containing scaffolds at day 7. At day 7, heparin-containing scaffolds had greater expression of *COL1A2* compared to day 1 than any other group.

**Figure 4:**
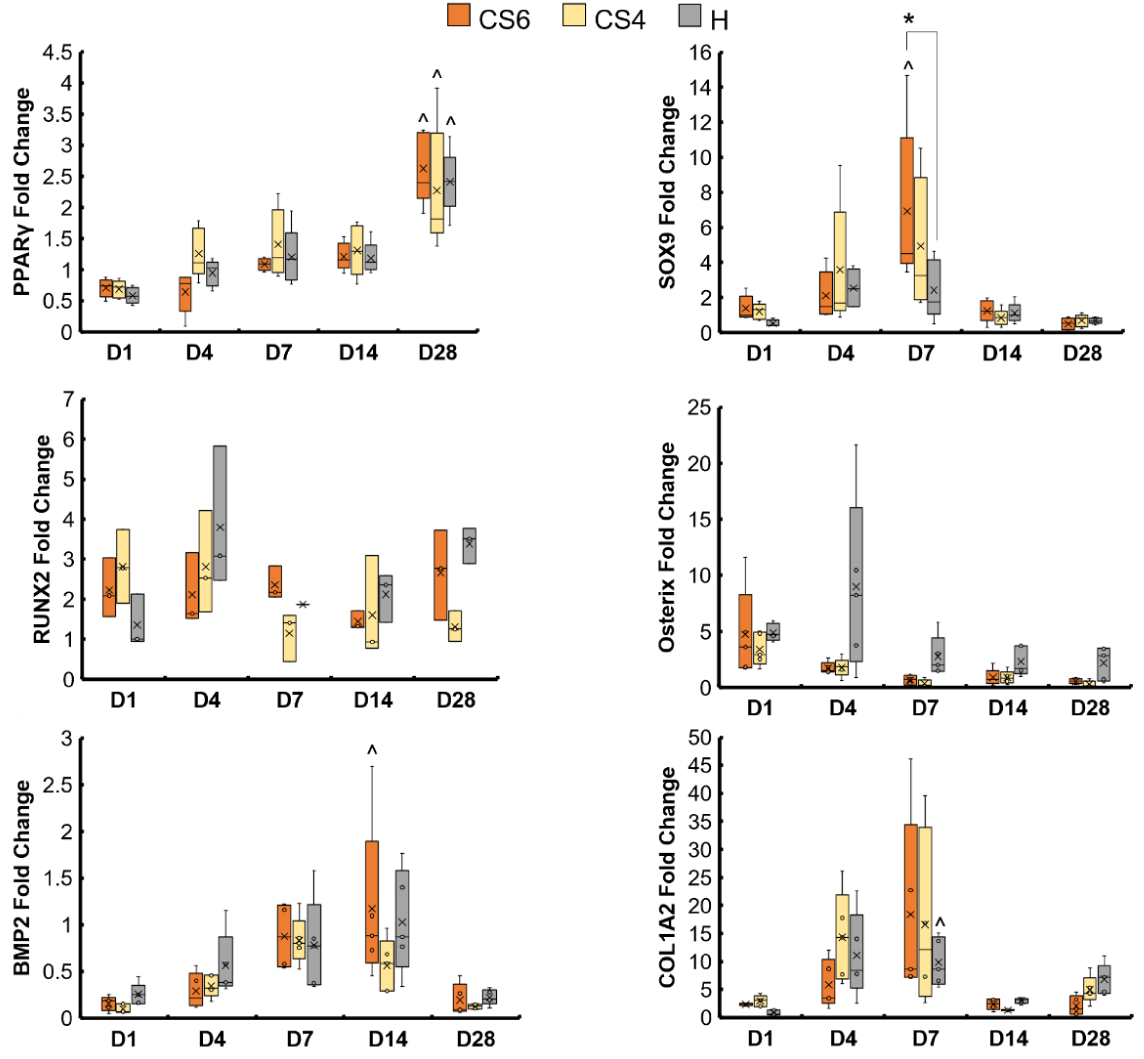
Osteogenic, chondrogenic, and adipogenic gene expression of mineralized collagen scaffolds with different GAGs, chondroitin-6-sulfate (CS6), chondroitin-4-sulfate (CS4), and heparin sulfate (H). * indicates one group is significantly (p < 0.05) greater than another group on the same day. ^ indicates one group was significantly (p < 0.05) greater than the same group on day 1. Data expressed as average ± standard deviation (n=5).

### 3.5. Greater mineral formation in anisotropic and CS6-containing scaffolds

Mineral formation in scaffolds was evaluated by Micro-CT and ICP analysis at day 28. As all scaffolds contained CaP mineral to begin with, results are reported as mineral content in scaffolds seeded with cells versus acellular scaffolds maintained in culture media. Micro-CT revealed anisotropic scaffolds had significantly greater mineral formation than isotropic scaffolds. Micro-CT revealed no differences in mineral formation between the different glycosaminoglycans, and all scaffolds had significantly greater mineral formation than day 0 unseeded scaffolds (**Fig. 5A, Supp. Fig. 4**). ICP analysis revealed that the CS6-containing scaffolds had significantly greater Calcium and Phosphorous than CS4- and heparin-containing scaffolds (**Fig. 5B**). Micro-CT was further validated by a greater Calcium and Phosphorous percent in the anisotropic variants.

**Figure 5:**
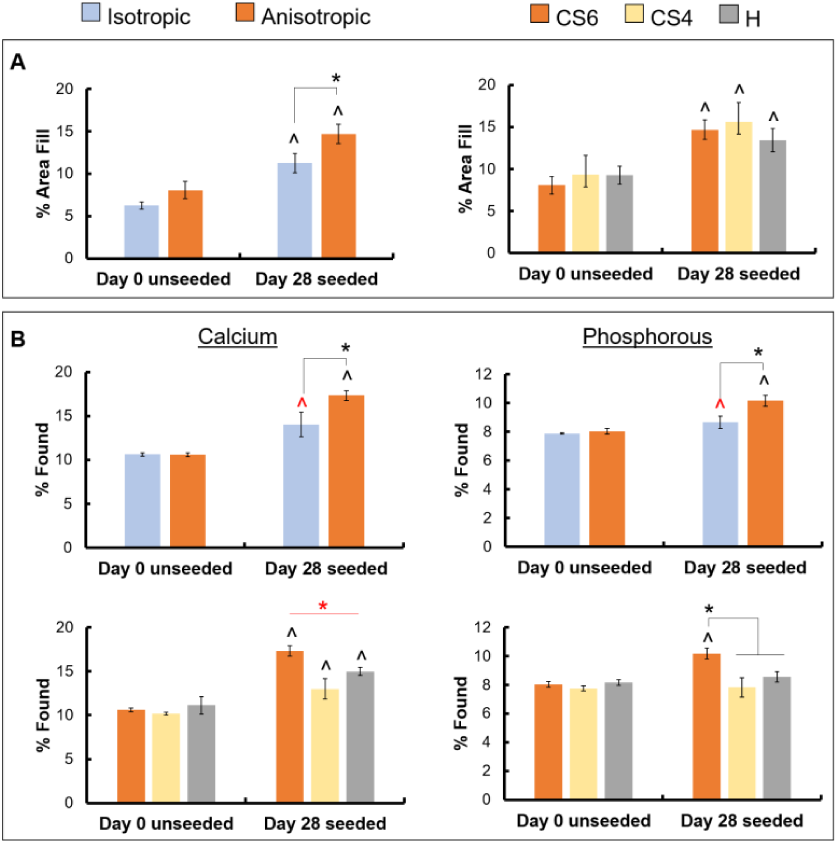
Mineral quantification of unseeded and hMSC seeded scaffolds by Micro-CT and ICP. (A) Micro-CT image stacks were processed in ImageJ with a threshold of 214/255 and the white area was compared to the total scaffold area to produce a % area fill. No statistical significance was found between the CS6, CS4, and H scaffold groups, however, all scaffolds had significantly (p < 0.05) greater % area fill at day 28 compared to day 0. * indicates there was a significant (p < 0.05) difference in % area fill between the isotropic and anisotropic groups seeded with cells (power: 0.78794) (B) ICP analysis of calcium and phosphorous content of scaffolds post-Micro-CT scanning and critical point drying compared to dry unseeded scaffolds. * indicates there is significantly (p < 0.05) greater % found Calcium and Phosphorous in the anisotropic and CS6 scaffold than all other scaffold types. ^ indicates significantly (p < 0.05) greater % found in at day 28 than day 0 of the same group. Data expressed as average ± standard deviation (n=3).

### 3.6. Greater osteogenic activity in scaffolds containing Heparin

Osteogenic activity was quantified by an Alkaline Phosphatase assay, again with MSC-seeded scaffolds compared to acellular controls. There was no difference in osteogenic activity between the isotropic and anisotropic scaffold variants, however, heparin-containing scaffolds had the greatest active bone formation day 28. (**Fig. 6, Table 3**). Interestingly, CS4-containing scaffolds had the lowest active bone formation at day 28.

**Table 3.**
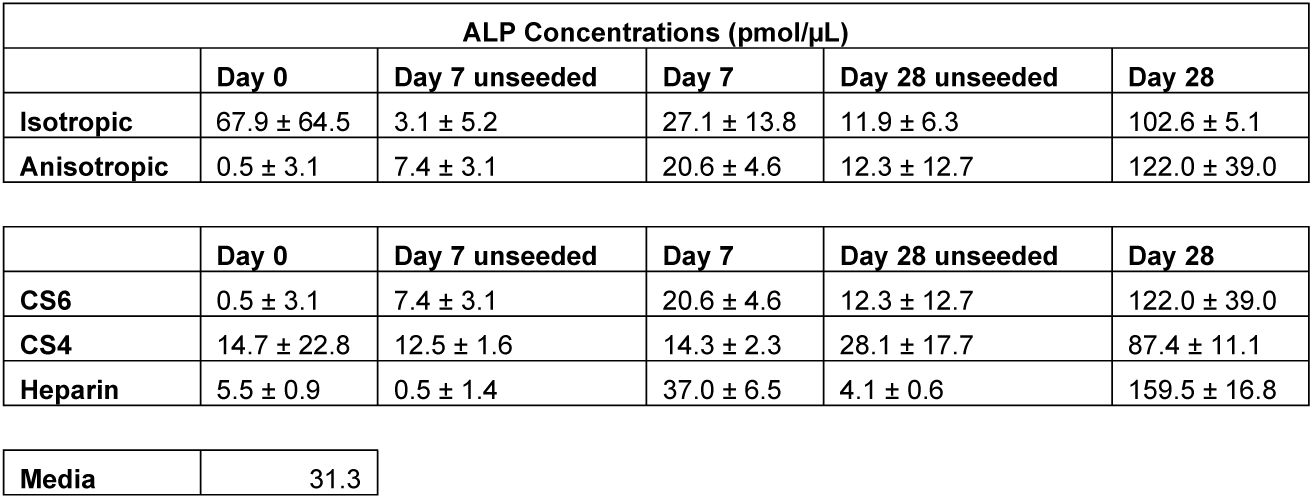
Alkaline phosphatase concentrations in unseeded and cell-seeded scaffolds with a media control.

| ALP Concentrations (pmol/μL) | | | | | |
| --- | --- | --- | --- | --- | --- |
|  | Day 0 | Day 7 unseeded | Day 7 | Day 28 unseeded | Day 28 |
| **Isotropic** | 67.9 ± 64.5 | 3.1 ± 5.2 | 27.1 ± 13.8 | 11.9 ± 6.3 | 102.6 ± 5.1 |
| **Anisotropic** | 0.5 ± 3.1 | 7.4 ± 3.1 | 20.6 ± 4.6 | 12.3 ± 12.7 | 122.0 ± 39.0 |

|  | Day 0 | Day 7 unseeded | Day 7 | Day 28 unseeded | Day 28 |
| --- | --- | --- | --- | --- | --- |
| **CS6** | 0.5 ± 3.1 | 7.4 ± 3.1 | 20.6 ± 4.6 | 12.3 ± 12.7 | 122.0 ± 39.0 |
| **CS4** | 14.7 ± 22.8 | 12.5 ± 1.6 | 14.3 ± 2.3 | 28.1 ± 17.7 | 87.4 ± 11.1 |
| **Heparin** | 5.5 ± 0.9 | 0.5 ± 1.4 | 37.0 ± 6.5 | 4.1 ± 0.6 | 159.5 ± 16.8 |

|  |  |
| --- | --- |
| **Media** | 31.3 |

**Figure 6:**
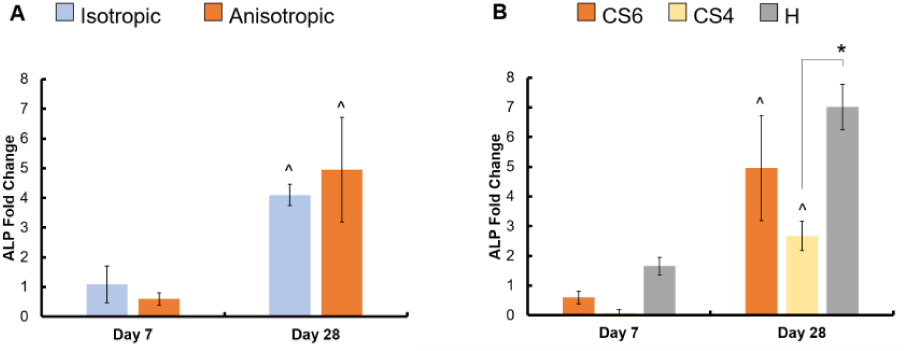
Active bone formation in scaffolds with varying pore orientation and glycosaminoglycan content. A colorimetric ALP assay was used to determine the ALP enzymatic activity in scaffolds after 7 and 28 days, which can be related to active bone formation. Values of the seeded scaffolds at days 7 and 28 were subtracted from unseeded scaffolds at the same days, then normalized to day 0 unseeded controls to achieve a fold change in ALP activity from the start of the experiment (1 represents activity at day 0). Chondroitin-6-sulfate, chondroitin-4-sulfate, and heparin sulfate-containing scaffolds are represented by CS6, CS4, and H (respectively). ^ indicates one scaffold group has significantly (p < 0.05) greater ALP activity at day 28 compared to the same group at day 7. ** indicates the heparin scaffold group had significantly (p < 0.05) greater ALP activity at day 7 compared to both CS6 and CS4. * indicates one scaffold group is significantly (p < 0.05) greater than a different group on the same day. Data expressed as average ± standard deviation (n=6).

### 3.7. No difference in migration efficiency between anisotropic and isotropic scaffolds containing CS6

Pore anisotropy did not increase cell invasion as measured via metabolic activity after 24hrs. Anisotropic scaffolds had a higher metabolic activity, however, this was not significantly greater than isotropic scaffolds (p = 0.37) (**Supp. Fig. 4**).

### 3.8. Secreteome from anisotropic and CS6-containing scaffolds showed increased levels of PDGF-BB and Angiogenin, respectively

PDGF-BB was significantly more expressed by cells maintained in anisotropic scaffolds for 7 days (**Table 4**). Angiogenin was significantly more expressed in CS6-containing scaffolds versus other GAG variants (Table 4). Overall, levels of anti-inflammatory cytokines remained the same, IL-6 and IL-8 pro-inflammatory cytokines had high fold changes. Notably, osteogenic cytokines OPN and OPG were upregulated in all scaffold variants but was not influenced by pore anisotropy or GAG content.

**Table 4.**
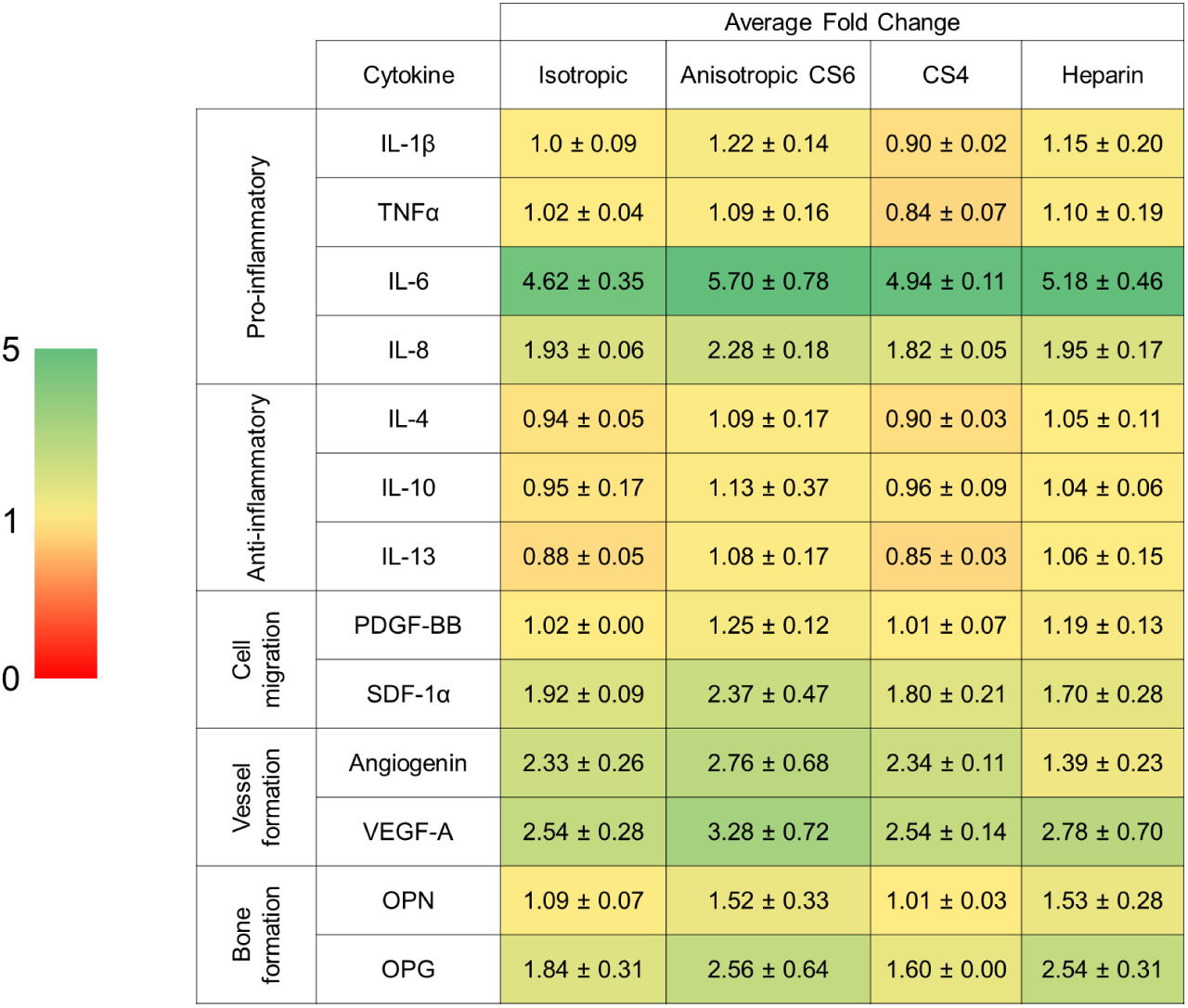
Secreted cytokines of hMSC seeded scaffolds with varying pore orientation. A cytokine assay was conducted on eluted media after 7 days from scaffolds with isotropic and anisotropic pores. Values of array intensity were measured with ImageJ, and a control array (exposed only to media) was used to determine fold change due to cellular activity. 5 key groups are shown – cytokines connected to inflammation, anti-inflammatory cytokines, cytokines connected to cell movement, angiogenic cytokines, bone formation cytokines. Green indicates cytokines with a higher fold change and yellow indicates limited or no change compared to media. * indicates a statistically significant (p < 0.05) increase in cytokine level for anisotropic pores over isotropic pores (power: 0.69176). Data expressed as average ± standard deviation (n=3).

## 4. Discussion

Craniomaxillofacial defects represent a unique and challenging type of bone defect. Herein, we explored modification of pore anisotropy and glycosaminoglycan content of a mineralized collagen scaffold under development for craniofacial bone repair. We hypothesized that anisotropic pores in the scaffold would enhance cell migration via directional guidance in order to subsequently improve osteogenic activity and new mineral biosynthesis. We further hypothesized that CS4 and Heparin in the scaffold would promote osteogenic activity and new mineral biosynthesis due to their potential to regenerate bone in literature ^33–36^.

We expected anisotropic pores to promote cell migration due to the aligned orientation of the pores; however, we saw no differences in either of these aspects between our anisotropic and isotropic pores. This contradicted some instances in literature, where accelerated migration could be achieved by anisotropic materials *in vivo* and *in vitro*, especially in aligned tissues ^53^. Bone is considered to be an aligned tissue, both the cortical and cancellous portions of bone having anisotropic behavior ^54^. It was of notice that a cytokine array revealed significantly greater expression of the chemokine PDGF-BB in anisotropic scaffolds than isotropic ones, thus a transwell study with longer timepoints may be of interest in the future to determine if migration effects are significant over longer periods of time. In addition, PDGF is important in the skeletal system, activating MSCs to differentiate to osteoblasts and activates macrophages ^55^. This could indicate that with significantly greater PDGF levels in anisotropic scaffolds, these scaffolds had greater MSC differentiation to osteoblasts. Although migration and activity of hMSCs was unaffected by anisotropy, this pore orientation change effected the ultimate behavior of the cells and mineral formation.

Both scaffold types had upregulated osteogenic genes *RUNX2, BMP2, Osterix*, and *COL1A2*, but anisotropic variants had a significant increase in chondrogenic gene, *SOX9*, at day 7. This difference in chondrogenic gene expression did not negatively impact osteogenic activity, as anisotropic scaffolds had significantly greater mineral formation than isotropic variants at the end of the study. *RUNX2* is a major transcription factor of bone, involved in the MSC transition to osteoblasts ^56^. There was no significant effect of pore orientation on *RUNX2* gene expression; however, this was upregulated throughout the study, indicating differentiation of hMSCs in both scaffold variants. The *COL1A2* gene is highly expressed in most connective tissues, such as bone, skin, and tendon ^57^. This gene was upregulated throughout the study, but expressed the greatest at day 7, indicating cells producing type I collagen, which could be related to bone formation occurring. *BMP2* is also involved in osteogenic differentiation, as well as bone formation ^58^. *BMP2* was only upregulated at days 7 and 14, possibly indicating that the greatest bone formation or differentiation was occurring at these timepoints. *Osterix* is important for late-stage bone formation, regulating the differentiation of osteoprogenitors to osteoblasts ^59^. *Osterix* was upregulated at the early-stages of the study, possibly indicating hMSCs were beginning to differentiate early-on. Interestingly, the expression of the osteoclast inhibitory factor OPG ^60^ was upregulated in all scaffold groups, particularly in the anisotropic mineralized scaffold. We have previously shown that mineralized collagen scaffolds containing isotropic pores and CS6 induce MSCs to produce OPG when seeded with MSCs to enhance bone regeneration ^44,45^. Suggesting the anisotropic mineralized scaffold may be particularly interesting for future studies to explore MSC-osteoclast interactions.

Although there were minimal differences in gene expression between isotropic and anisotropic scaffolds, we observed greater mineral formation via ICP and Micro-CT analyses in anisotropic scaffolds. At the end of the study, total amounts of Calcium and Phosphorous were greater in the anisotropic scaffold compared to the isotropic scaffold. Interestingly, a study by Bobbert *et al*. revealed that aligned fibers in PCL structures had a lower ALP activity than random fibers ^61^. This difference in results could be attributed to a difference in metabolic activity of PCL structures; however, our scaffolds had no metabolic activity differences, indicating no bias on ALP activity. Of note, another study by Seong *et al*. indicated that small channels enhanced proliferation and migration *in vitro* and pore sizes of 270μm were best for cell migration *in vivo* ^62^. This could explain the lack of migration and metabolic activity differences in our scaffolds, which had average pore diameters in the range of 150 – 200 μm.

We expected addition of chondroitin-4-sulfate or heparin sulfate to increase osteogenic activity and bone formation versus chondroitin-6-sulfate-containing scaffolds. This was based on observations in the literature that CS4 is more involved in the bone formation process while CS6 is involved in cartilage formation ^33^. We found no differences in metabolic activity between any of the glycosaminoglycans, but all promoted a greater than 2.5-fold change in activity after 28 days. Overall, osteogenic activity was observed throughout all variants. However, heparin-functionalized variants increased relative expression (vs. day 1) of *COL1A2* as well as greater expression of *Osterix* than CS4 or CS6 variants, suggesting more osteoblastic differentiation in heparin scaffolds at this period. CS6-functionalized variants also showed an early (day 7) spike in *SOX9* chondrogenic gene expression. However, overall, all osteogenic genes were upregulated for most of the study regardless of glycosaminoglycan type, suggesting future studies to look at long-term functional responses *in vitro* and *in vivo*.

Analysis of mineral formation via Micro-CT found increased mineral deposition in all variants but was unable to resolve difference in mineral formation at 28 days in culture. Analysis of ALP levels indicated that active bone formation was greatest in Heparin scaffolds at day 7 and 28, although, there were no differences in ALP activity between Heparin and CS6 scaffolds at day 28. Overall, CS4-containing scaffolds showed the smallest amount of mineral formation compared to CS6 or Heparin scaffolds. While not examined explicitly here, strategies to improve angiogenesis are essential for large volume bone regeneration; analysis of the secretome of MSCs in all scaffolds revealed CS6 or Heparin-containing anisotropic scaffolds showed significantly greater angiogenin expression than CS4 variants. Overall, our data suggest anisotropic mineralized collagen scaffolds containing either CS6 or Heparin GAGs have the greatest potential for bone regeneration.

Ongoing efforts are seeking to expand on our library of anisotropic scaffolds to explore the role of anisotropy and pore size on cell migration. Observations from the secretome analysis also suggest the potential for examining the role of scaffold anisotropy and GAG composition on inflammatory response. Although CS6 and Heparin proved to have the most potential for bone regeneration, the immune response and vessel formation are still two important factors for bone repair. The cytokine array also revealed that there was a high fold change of pro-inflammatory IL-6 and IL-8 cytokines in all glycosaminoglycans examined. Heparin induces a more pro-inflammatory phenotype and activates the NF-κB pathway, which may play a role in inflammatory diseases, whereas chondroitin sulfates do not ^63–65^. In addition, CS4 has been shown to inhibit reactive oxygen species and decrease pro-inflammatory factors such as IL-1β and TNFα ^66^. Although larger pores of 270μm may have been shown to promote cell migration, pores on the order of 40μm may prove more beneficial to avoiding a negative immune response and fibrosis ^67–69^. Future work will involve examining the effect of these different GAGs in mineralized collagen scaffolds on the immune response, as bone formation may be improved by CS6 or Heparin, but each GAG may induce a different immune response. We also intend to expand our analysis of angiogenic responses in anisotropic mineralized collagen scaffolds, building on recent observation that vessel formation can depend on fiber alignment ^61^.

## 5. Conclusions

This study examined the role of pore anisotropy and GAG composition on MSC osteogenic activity in a class of mineralized collagen scaffolds under development for CMF bone repair applications. We hypothesized that anisotropic scaffolds would promote greater cell migration, osteogenic activity, and bone formation. We found that anisotropic pores did not increase cell activity or migration but did improve overall bone formation. We also hypothesized that inclusion of chondroitin-4-sulfate and heparin sulfate would increase cell activity and bone formation in mineralized collagen scaffolds. Notably, chondroitin-6-sulfate and heparin sulfate had the greatest effect on improved mineral formation. These efforts represent an important step toward optimizing mineralized collagen scaffolds for bone regeneration. These studies also set the stage to examine the role of anisotropy and GAG content on inflammatory and angiogenic responses as well as crosstalk between these axes and MSCs.

## Acknowledgements

This work was supported by the Office of the Assistant Secretary of Defense for Health Affairs Broad Agency Announcement for Extramural Medical Research through the Award No. W81XWH-16-1-0566. Opinions, interpretations, conclusions and recommendations are those of the authors and are not necessarily endorsed by the Department of Defense. Research reported in this publication was also supported by the National Institute of Dental and Craniofacial Research of the National Institutes of Health under Award Number R21 DE026582. The content is solely the responsibility of the authors and does not necessarily represent the official views of the NIH. We are grateful for the funding for this study provided by the NSF Graduate Research Fellowship DGE-1144245 (MJD).

The authors would like to acknowledge the University of Illinois Roy J. Carver Biotechnology Center for assistance with real-time PCR. The authors also acknowledge assistance from the School of Chemical Science Microanalysis Lab. This research was carried out in part at the Imaging Technology Group within the Beckman Institute for Advanced Science and Technology at the University of Illinois at Urbana-Champaign. The authors would like to thank Leilei Yin for help with microcomputed tomography, as well as Scott Robinson and Cate Wallace for assistance with critical point drying and scanning electron microscopy. Additional support was provided by the Chemical and Biomolecular Engineering Dept. and the Carl R. Woese Institute for Genomic Biology (BACH) at the University of Illinois at Urbana-Champaign.

---

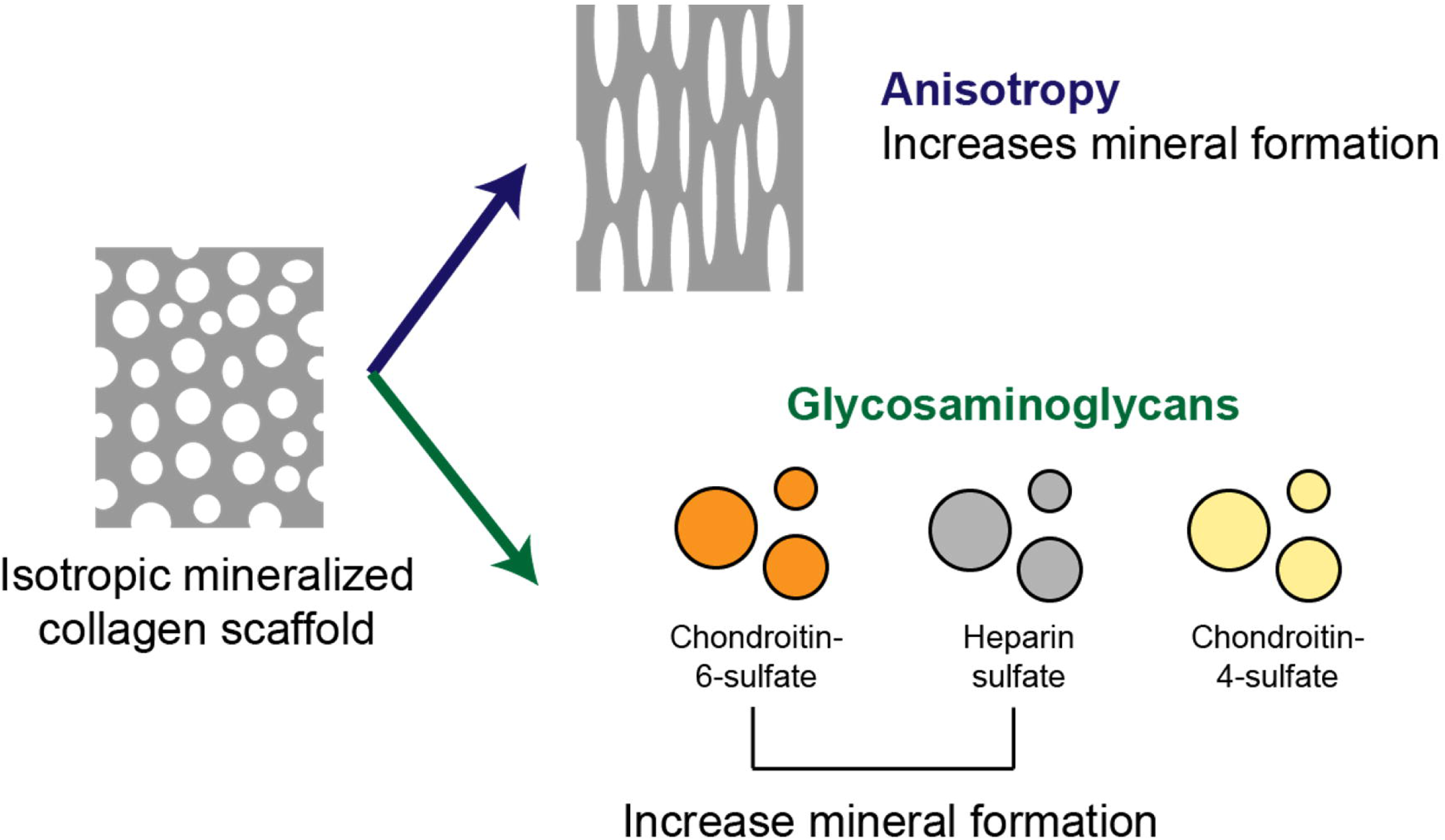

Mineralized collagen scaffolds were modified to include anisotropic pore architecture and one of three glycosaminoglycans in order to improve bone mineral formation *in vitro*.

