## Supplemental Data for "Anisotropic mineralized collagen scaffolds accelerate osteogenic response in a glycosaminoglycan-dependent fashion"

### Influence of pore anisotropy and glycosaminoglycan content on osteogenic activity of mineralized collagen scaffolds

[1] Dept. of Materials Science and Engineering
[2] School of Chemical Sciences
[3] Dept. Chemical and Biomolecular Engineering
[4] Carl R. Woese Institute for Genomic Biology
University of Illinois at Urbana-Champaign
Urbana, IL 61801

### Supplementary Figures

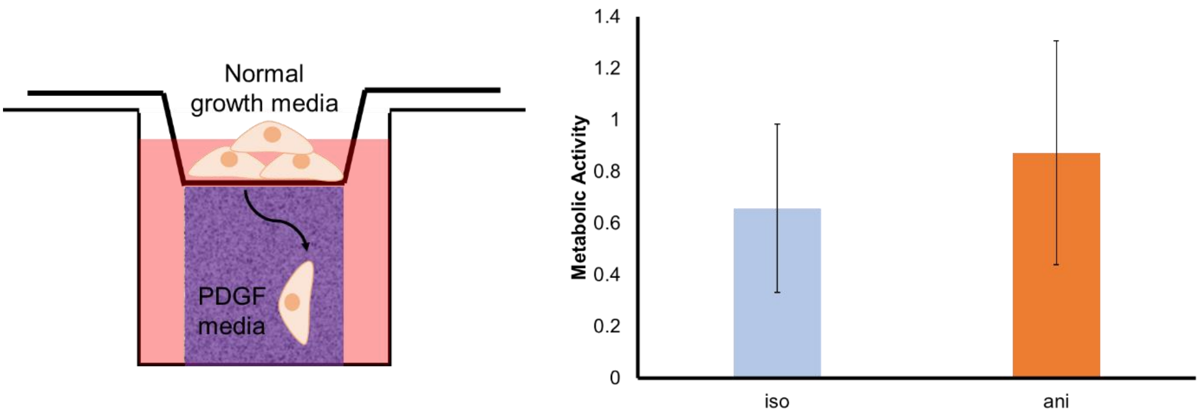


***Supplementary Figure 1:*** Representative image of transwell-seeded scaffolds and the metabolic activity of isotropic and anisotropic scaffold after 24-hours. Anisotropic and isotropic scaffolds containing chondroitin-6-sulfate were placed below transwells in media containing PDGF growth factor and 50,000 hMSCs were seeded on transwell membrane above scaffolds. Cells were allowed to migrate into scaffolds for 24 hours. After 24 hours, scaffolds were removed and the metabolic activity of scaffolds was measured with an alamar blue assay. A value of 1 represents the metabolic activity of 50,000 cells seeded on the transwells as the start of the experiment. Data expressed as average ± standard deviation (n=6).

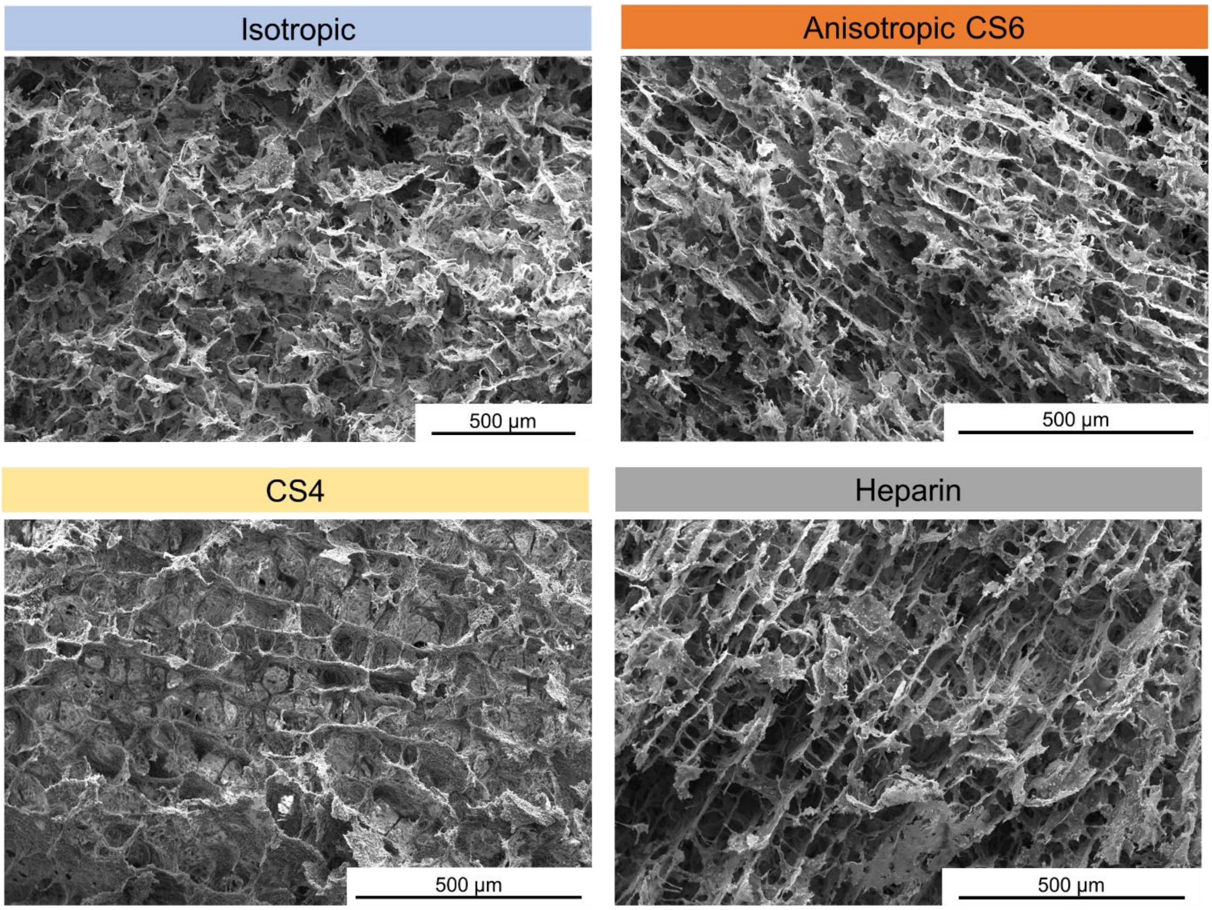


**Supplementary Figure 2.** SEM of pore structure of isotropic chondroitin-6-sulfate (CS6), anisotropic chondroitin-6-sulfate (CS6), anisotropic chondroitin-4-sulfate (CS4), and anisotropic heparin sulfate (Heparin) scaffolds. Isotropic scaffolds demonstrate random round-shaped pore structure and anisotropic scaffolds demonstrate square, elongated pores regardless of glycosaminoglycan content.

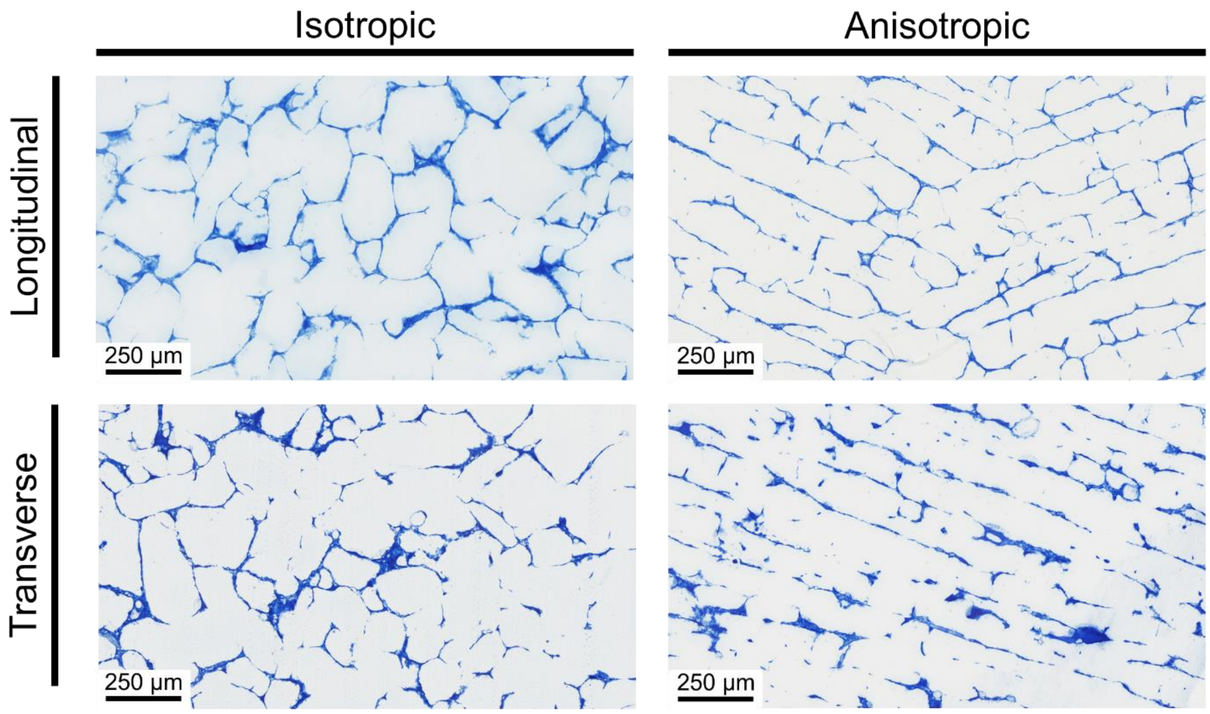


**Supplementary Figure 3. Representative aniline blue stained sections of isotropic and anisotropic mineralized collagen scaffolds with glycosaminoglycan chondroitin-6-sulfate.**

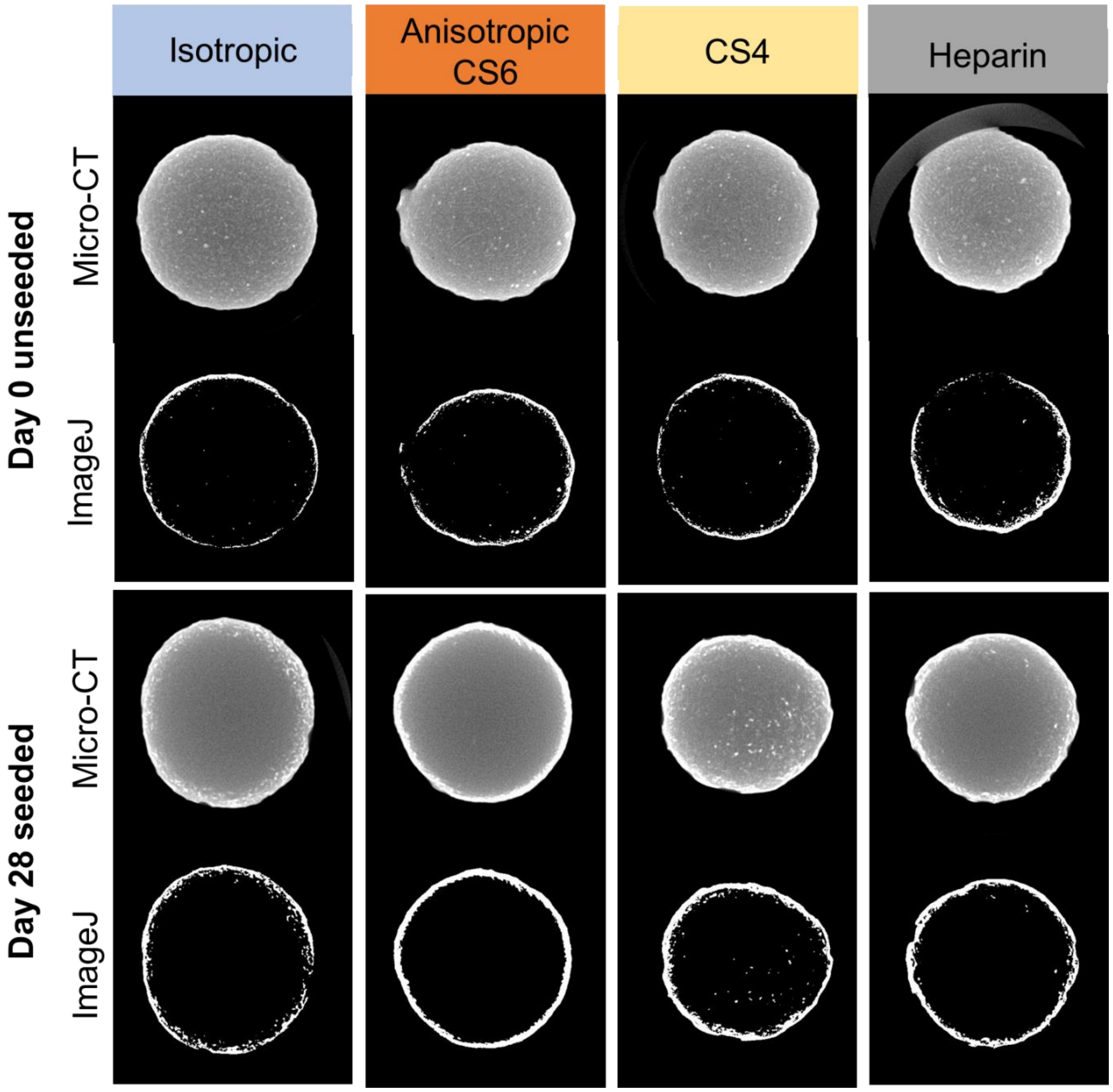


**Supplementary Figure 4. Representative Micro-CT scans and their ImageJ threshold equivalents.** Chondroitin-6-sulfate, chondroitin-4-sulfate, and heparin sulfate-containing scaffolds are represented by CS6, CS4, and H (respectively). The scaffolds seeded with hMSCs for 28 days are compared to unseeded scaffolds after the hydration process. Quantitative graphs were produced by comparing the scaffold area with the ImageJ white areas in a stack of Micro-CT images.

### Supplementary Tables

**Supplementary Table 1. Parameters for microwave assisted acid digestion of samples.**

| Digestion Parameters | Values |
|---|---|
| Power | 1800 W |
| Temperature | 200 $^0$C |
| Ramp Time | 30 minutes |
| Hold Time | 20 minutes |

**Supplementary Table 2. Parameters for analysis of calcium and phosphorus via Optima 8300 ICP-OES.**

| ICP-OES Parameters | Values |
|---|---|
| RF Power | 1500 Watts |
| Nebulizer | GemCone Low Flow |
| Nebulizer Gas Flow rate | 0.85L/min |
| Plasma Gas Flow rate- Argon | 10L/min |
| Sample Flow rate | 1.50mL/min |

The following emission lines were used for the quantification of Ca and P
Ca (II)-319.93 nm
P (I)- 213.62 nm
